# Characterization and utility of novel monoclonal antibodies to cholera toxin B subunit

**DOI:** 10.1101/2022.10.25.513727

**Authors:** Noel Verjan Garcia, Ian Carlosalberto Santisteban Celis, Matthew Dent, Nobuyuki Matoba

**Affiliations:** UofL Health – Brown Cancer Center, University of Louisville School of Medicine, Louisville, KY, USA; Center for Predictive Medicine, University of Louisville School of Medicine, Louisville, KY, USA; Department of Pharmacology and Toxicology, University of Louisville School of Medicine, Louisville, KY, USA

**Keywords:** cholera toxin B subunit, EPICERTIN, monoclonal antibody, hybridoma, surface plasmon resonance, flow cytometry, immunohistochemistry

## Abstract

Cholera toxin B subunit (CTB) is a potent immunomodulator exploitable in mucosal vaccine and immunotherapeutic development. To aid in the characterization of pleiotropic biological functions of CTB and its variants, we generated a panel of anti-CTB monoclonal antibodies (mAbs). By ELISA and surface plasmon resonance, two mAbs, 7A12B3 and 9F9C7, were analyzed for their binding affinities to cholera holotoxin (CTX), CTB, and EPICERTIN: a recombinant CTB variant possessing mucosal healing activity. Both 7A12B3 and 9F9C7 bound efficiently to CTX, CTB, and EPICERTIN with equilibrium dissociation constants at low to sub-nanomolar concentrations but bound weakly, if at all, to *Escherichia coli* heat-labile enterotoxin B subunit. In a cyclic adenosine monophosphate (cAMP) assay using Caco2 human colon epithelial cells, the 7A12B3 mAb was found to be a potent inhibitor of CTX, whereas 9F9C7 had relatively weak inhibitory activity. Meanwhile, the 9F9C7 mAb effectively detected CTB and EPICERTIN bound to the surface of Caco2 cells and mouse spleen leukocytes by flow cytometry. Using 9F9C7 in immunohistochemistry, we confirmed the preferential localization of EPICERTIN in colon crypts following oral administration of the protein in mice. Collectively, these mAbs provide valuable tools to investigate the biological functions and preclinical development of CTB variants.

## INTRODUCTION

Cholera toxin B subunit (CTB, approx. 58 kDa), is a nontoxic component of the cholera toxin (CTX) released by *Vibrio cholerae*, a Gram-negative bacterium causing profuse secretory diarrhea.^1^ CTB is a potent immunomodulator consisting of five identical polypeptide chains non-covalently associated in a ring-shaped pentameric structure that mediates CTX binding to the monosialoganglioside GM 1 receptor.^2–4^ CTB has been produced in various recombinant expression platforms as model vaccines,^5–8^ as it induces a strong immune response characterized by neutralizing antibodies to the CTX via oral administration.^9,10^ CTB has also been used as a molecular scaffold for immunization^11,12^ and for induction of peripheral (oral) immunological tolerance to suppress allergies and various autoimmune disorders such as experimental autoimmune encephalitis (EAE), diabetes, arthritis and uveitis in an antigen-specific manner.^13–17^ CTB is considered a mucosal immunomodulator that induces a strong mucosal IgA response while it seems to suppress systemic T helper (T_H_1, T_H_2 and T_H_17) responses through the induction of interleukin-10 (IL-10) and transforming growth factor-β (TGF-β)-mediated regulatory T (Treg) induction and suppression of proinflammatory IL-6,^18^ although the exact mechanism behind such immunomodulation remains elusive.

The pleiotropic functions of CTB have recently been expanded through the construction of CTB variants with novel biological activities, including a plant-made CTB variant modified with C-terminal hexapeptide extension containing a KDEL endoplasmic reticulum (ER) retention motif,^19^ later designated as EPICERTIN,^20^ which showed a significant enhancement of mucosal wound healing activity in the colon.^21–25^ In mice, oral administration of EPICERTIN increased specific subsets of immune cell populations, such as macrophages, natural killer cells, Treg cells and T_H_17 cells in the colon lamina propria and upregulated wound healing pathways mediated by TGF-β in the colon. Mucosal healing effects of EPICERIN were confirmed in an *in vivo* dextran sodium sulfate (DSS) acute colitis model, an azoxymethane/DSS model of ulcerative colitis and tumorigenicity,^21^ and a model of chronic colitis-like colon inflammation induced by repeated doses of DSS, where the wound healing effects were not neutralized by the induction of mucosal anti-CTB antibodies.^24^ The prolonged residence of EPICERTIN in the ER, through interaction with the KDEL receptor, appears to trigger an unfolded protein response, which leads to the activation of TGF-β signaling and wound healing pathways in colon epithelial cells.^23^ However, the mucosal healing effects could also originate from modulation of mucosal immune cells. Indeed, early studies reported multiples roles of CTX and CTB in mouse and human leukocytes. For example, CTB inhibited T-cell proliferation induced by mitogens (e.g. Concanavalin A) and antigens,^26^ and antigen-conjugated CTB promoted its presentation and lowered the threshold of antigen required for T cell activation.^27^ Therefore, to facilitate the investigation of the mechanisms behind the biological activities of EPICERTIN and other CTB-derived molecules, we aimed at generating monoclonal antibodies (mAbs) against CTB that aid in the characterization of those proteins.

Here, we report on the isolation and characterization of two novel mAbs generated against CTB, demonstrating their unique and distinct binding profiles by means of ELISA, surface plasmon resonance (SPR) and cyclic adenosine monophosphate (cAMP) assays. In addition, we show the utility of one of the newly isolated mAbs in detecting CTB bound both to murine leukocytes by flow cytometry and to colon tissues by immunohistochemistry (IHC), which will aid in investigating as-yet-uncharacterized biological mechanisms of action of CTB and its derivatives as candidate mucosal vaccines and immunotherapeutics.

## METHODS

### Animal care

Immunization of Wistar rats for hybridoma generation was conducted by GenScript USA, Inc. (Piscataway, NJ), whereas flow cytometry and IHC experiments using C57BL/6J mice were performed at the University of Louisville. Studies were approved by each institution’s Institutional Animal Care and Use Committee. General procedures for animal care and housing in these studies were in accordance with the current Association for Assessment and Accreditation of Laboratory Animal Care recommendations, current requirements stated in the Guide for the Care and Use of Laboratory Animals (National Research Council), and current requirements as stated by the U.S. Department of Agriculture through the Animal Welfare Act and Animal Welfare regulations (July 2020).

#### Reagents

Cholera enterotoxin (C8052), cholera toxin B-subunit (CTB; C9903) and heat labile enterotoxin B subunit (LTB; E9656) of *Escherichia coli* were purchased from Sigma (Sigma Aldrich). PhenoVue Fluor 594-WGA was obtained from PerkinElmer (Perkin Elmer Health Science Inc. Boston MA, USA), and Alexa Fluor 647-anti mouse/human CD324 (E-cadherin) (clone DECAMA-1) was obtained from Biolegend.

#### Generation of anti-CTB mAbs

To facilitate the generation of CTX-neutralizing mAbs, an Asn4-glycosylated CTB variant produced in *Nicotiana benthamiana* ^19,29^ (gCTB) was used as an antigen for immunization. The plant-expressed gCTB was purified by metal-affinity chromatography followed by CHT hydroxyapatite chromatography, as described previously ^19^. The purified protein was used to immunize Wistar rats to generate a panel of monoclonal antibodies by Genscript USA, Inc. using standard hybridoma technology ^30,31^. Briefly, Wistar rats were immunized with 50 μg of keyhole limpet hemocyanin conjugated gCTB (gCTB-KLH) and subsequently boosted with 25 μg gCTB-KLH every two weeks, for a total of three booster doses. Two weeks after the last booster immunization, the lymphocyte population from spleens were isolated and used to generate immortalized hybridomas by fusion with the Sp2/0-Ag14 myeloma cells following standard methods. Hybridomas were grown in Hypoxanthine-aminopterin-thymidine medium supplemented with IL-6, and the culture supernatants were screened for Abs reactive with CTB by direct ELISA and GM1-capture ELISA. Two anti-CTB mAbs-producing hybridomas named 7A12B3 and 9F9C7 were selected and expanded, and mAbs were produced in hybridoma cell culture supernatants using the invitro roller bottle cell culture method followed by protein G-affinity chromatography purification. They were both determined to be of the IgG2a subclass by using the Pro-Detect^TM^ Rapid Antibody Isotyping Kit-Rat (Thermo Fisher Scientific).

#### Enzyme-linked immunosorbent assays

Binding affinities of 7A12B3 and 9F9C7 mAbs to CT were determined by antigen (CT, CTB, LTB)-capture enzyme-linked immunosorbent assays (ELISA) and competitive GM1/KDEL-ELISA as described previously^22^ with minor modifications. Briefly, 96-well cell culture plates (Nunc, MaxiSorp) were coated with 300 ng/mL of CTB, CT, or LTB in 50 mM carbonate coating buffer (NaCO_3_, NaHCO_3_, 0,1% NaN_3_) or 2 μg/mL GM-1 ganglioside (Sigma, St. Louis, Mo) at 4 °C overnight or at room temperature for 2 hr, respectively. The plates were washed 3 × with PBST (phosphate buffered saline + 0.05% Tween 20) and blocked with 5% non-fat dry milk in PBST for 1 hr at room temperature or left overnight at 4 °C. Serial dilutions of hybridoma supernatants (1: 100 dilution) or 7A12B3 and 9F9C7 mAbs or the recombinant proteins and the standards in PBST containing 1 % non-fat dry milk were added after 3 × washes with PBST and incubated at 37 °C for 1 hr. After 3 washes with PBST, a horseradish peroxidase (HRP)-conjugated anti-mouse IgG secondary antibody (multi-species SP ads-HRP-conjugated goat anti-mouse IgG (H+L) antiserum, Southern Biotech, Birmingham, AL, USA) diluted at 1:100.000 was added and incubated for 1 hr at room temperature. The GM1/KDEL-ELISA plates were incubated with a primary mouse anti-KDEL mAb (10C3, Enzo Life Sciences; 1:1000 diluted) at 37 °C for 1 hr followed by the multi-species SP ads-HRP-conjugated goat anti-mouse IgG (H+L) antiserum (Southern Biotech) diluted at 1:20.000 and incubated at 37 °C for 1 hr. The plates were washed 3 × and the horseradish peroxidase substrate tetramethylbenzidine, (TMB, 100 μL/well) was added and incubated for 2 min at room temperature before stopping the reaction with 100 μl/well of 0.6 N H_2_SO_4_ stop solution. Absorbance at 450 nm was measured using a BioTek Synergy HT microplate reader (Winooski, VT, USA).

#### Surface plasmon resonance

Binding affinities of 7A12B3 and 9F9C7 mAbs to CT, commercial CTB, and EPICERTIN were also determined by surface plasmon resonance (SPR) using the Biacore Gold Seal T200 (GE Healthcare) equipped with a CM5 sensor chip as previously described ^19^. The ligands 7A12B3 (150 kDa) and 9F9C7 (150 kDa) mAbs were immobilized on the carboxylated dextran matrix of a CM5 chip sensor surface using amine-coupling chemistry. The surfaces of flow cells were activated with a 1:1 mixture of 0.1 M NHS (N-hydroxysuccinimide) and 0.4 M EDC (3-(N,N-dimethylamino) propyl-N-ethylcarbodiimide) at a flow rate of 5 μl/min for 14 min. The ligands at a concentration of 5 μg/ml in 10 mM sodium acetate, pH 4.0, were immobilized at a density of 205-220 RU (7A12B3) and 709 (9F9C7). Flow cells 1 and 2 were left as a reference blank. Both surfaces were blocked with 1 M ethanolamine, pH 8.0, with a 7 min injection time. Running buffer was 10 mM HEPES, 150 mM NaCl, 0.005% P20, pH 7.4. To collect steady state data and kinetic binding of analytes CT, CTB, and EPICERTIN to immobilized 7A12B3 mAb, the analytes were diluted to 26.9 nM (1.6 μg/mL) in running buffer. For 9F9C7 the analytes CT, CTB, and EPICERTIN were diluted to 1670 nM (100 μg/mL) in running buffer. Samples were 2-fold serially diluted and injected at a flow rate of 10 μL/min and 30 μL/min at 25°C, respectively. The complexes were allowed to associate and dissociate for 600 s and 900 s, respectively. The surfaces were regenerated with 10 mM Glycine pH 2.0 for 60 s. Triplicate injections (in random order) of each sample and a buffer blank were flowed over the two surfaces. Data were collected at a rate of 1 Hz. The data were fit to a simple 1:1 interaction model using the global data analysis option available within Biacore Evaluation software.

#### Intracellular cAMP in Caco2 cell line

Caco2 cells were grown in 6-wells and 12-wells cluster plates (Thermo Scientific Nunc Cell-Culture Treated, Roskilde, Denmark) at a density of 7×10^5^ Caco2 cells/well containing EMEM (Gibco BRL) medium supplemented with 20% FBS, 5 mM HEPES, 5 mM NEAA, 5 mM Sodium Pyruvate and Penicillin/Streptomycin for 24 hr. The culture medium was removed, and the cells were washed twice with PBS and then incubated in serum-free EMEM medium containing 2.5 mM HEPES, 0.01% bovine serum albumin, 1 mM 3-Isobutyl-1-methylxanthine (MP Biomedicals, LLC, Solon, Ohio), and the indicated concentrations of 7A12B3 and 9F9C7 mAbs, and 0.5 μg/mL (5.88 pmoles) of CT (Sigma, St. Louis, Mo) at 4 °C (on ice) or rat IgG isotype control for 30 min. The plates were subsequently transferred to 37 °C in a 5% CO_2_ incubator for 2 hr. The culture plates were centrifuged at 1000 × g, the culture medium removed, and the cells washed twice with PBS before incubation in 200 μL of 0.1 N HCl for 20 min at room temperature to allow cell lysis and cAMP extraction^32^. cAMP was detected using a sensitive colorimetric ELISA-based kit (Enzo Life Sciences, Farmingdale, NY) following the manufacturer’s conditions. A standard curve was constructed with known concentrations of cAMP and the cAMP levels in the samples (pmol/7×10^5^ Caco2 cells) were determined according to the equation generated by the standard curve.

#### Cell isolation and flow cytometry

C57BL/6J female mice were obtained from Jackson Laboratories (Bar Harbor, ME) and used between 8-12-week-old. The spleens of naïve mice were collected and minced, and the cell suspension passed through a 40 μm cell strainer after treatment with ACK buffer to lyse red cells. The cells were counted, the Fc receptors were blocked with mouse γ-globulins (20 μg/mL), and the cells were subsequently incubated with EPICERTIN or an EPICERTIN variant with Glv3%→Asp mutation (EPICERTIN^G33D^;^33^ 5 μg/mL) for 30 min on ice. After two washes with FACS buffer, unlabeled 9F9C7 mAb was added at 5 μg/mL and the cell suspension incubated on ice for 30 min. Later a FITC-conjugated goat anti-Rat IgG (Poly4054) antibody was added together with fluorochrome-labeled antibodies to cell specific markers, including PE-conjugated anti-CD4 (RM4-5), PE-conjugated anti-IA-IE (M5/114.15.6), PE-conjugated anti-Ly6C (Hk1.4), PE-conjugated anti-Gr-1 (RB6-8C5), PE or APC-conjugated anti-CD11c (N418), APC-conjugated anti-CD8a (53-6.7), APC-conjugated anti-CD11b (M1/70), APC-conjugated anti-Ly6G (1A8), APC-conjugated F4/80 (BM8) and PE or APC-conjugated Rat IgG2a κ isotype control (RTK2758) antibodies, all from Biolegend. APC-conjugated anti-CD19 (1D3) was from eBioscience. Flow cytometric analysis was performed on a FACSCalibur or a BD SLRFortessa (BD Biosciences) and the data were processed with FlowJo_v10.8.0_CL software. The geometric mean fluorescence intensity values generated by the goat-anti Rat IgG secondary antibody were subtracted from that of 9F9C7 anti-CTB specific mAb. The procedures with mice were approved by the Institutional Animal Care and Use Committee of University of Louisville.

#### Immunohistochemistry

EPICERTIN in PBS (3 μg/100μL) was administered by oral gavage to C57BL/6J mice after neutralization of the gastric acid with 200 μL of sodium bicarbonate (30 mg/mL). The mice were sacrificed at 0, 3, 6, 12 and 24 hr, and the colon tissues were washed with PBS and embedded in OCT compound to make 0.7 μm thick frozen sections. Cryosections of the colon tissue were fixed in 100% Methanol at −20 °C for 3 min, dried and blocked with 10% FBS in PBS containing 20 μg/mL mouse γ-globulins for 1 hr at RT. The tissue sections were stained with 5 μg/mL Pheno Vue Fluor 594-WGA (PerkinElmer Health Sciences), 2 μg/mL anti-E-cadherin, and 2 μg/mL anti-CTB (9F9C7 mAb) for 1 hr at RT. The slides were washed in PBS, images were collected with a Nikon A1R Confocal laser scanning microscope using 20 × and 60 × magnification lenses with appropriate channels, and the data were processed with the NIS Elements imaging software.

#### Data analysis

The half-maximal effective concentration (EC_50_) values were determined by non-linear regression analysis using Prism c.9.1.0. One-way ANOVA with Bonferroni’s multiple comparison posttest was used to analyze absorbance values of cAMP levels, using Prism v.9.1.0 (GraphPad Software, La Jolla, CA, USA). A value of *p* < 0.05 was considered significant.

## RESULTS

### Generation of anti-CTB mAbs

Three Wistar rats were immunized with a KLH-conjugated gCTB. All rats consistently showed high anti-CTB serum antibody titers after three doses of the antigen, as analyzed by both CTB and GM1-capture direct ELISA. Given that rat #1 showed an optimal response ratio for direct vs. GM1-bound CTB at a higher dilution factor (1:243,000), indicative of a significant proportion of antibodies targeting the GM1-binding facet of CTB, this rat was selected for hybridoma generation. A total of 40 positive hybridoma clones were obtained. Hybridoma supernatants were analyzed for the presence of anti-CTB antibodies by GM1- and CTB-capture ELISAs (**Figure 1A**). At this stage, the 7A12, 8F8 and 9F9 hybridomas were initially selected based on the high binding affinity to immobilized CTB in direct antigen-capture ELISA and in GM1-capture ELISA. Of note, among the three hybridomas, 7A12 and 9F9 mAbs showed very distinctive CTB-binding patterns in ELISA (**Figure 1B**), where the binding signal of 7A12 mAb was almost completely abolished in GM1-capture ELISA whereas 9F9 mAb showed similar binding responses regardless of the ELISA formats. Thus, we proceeded with subculturing of these two hybridomas under limiting dilution conditions to isolate single clones, and the resultant 7A12B3 and 9F9C7 hybridomas were selected for subsequent studies. DNA sequence analysis revealed that both 7A12B3 and 9F9C7 mAbs are of the IgG2a isotype with kappa light chains (data not shown).

**Figure 1.**
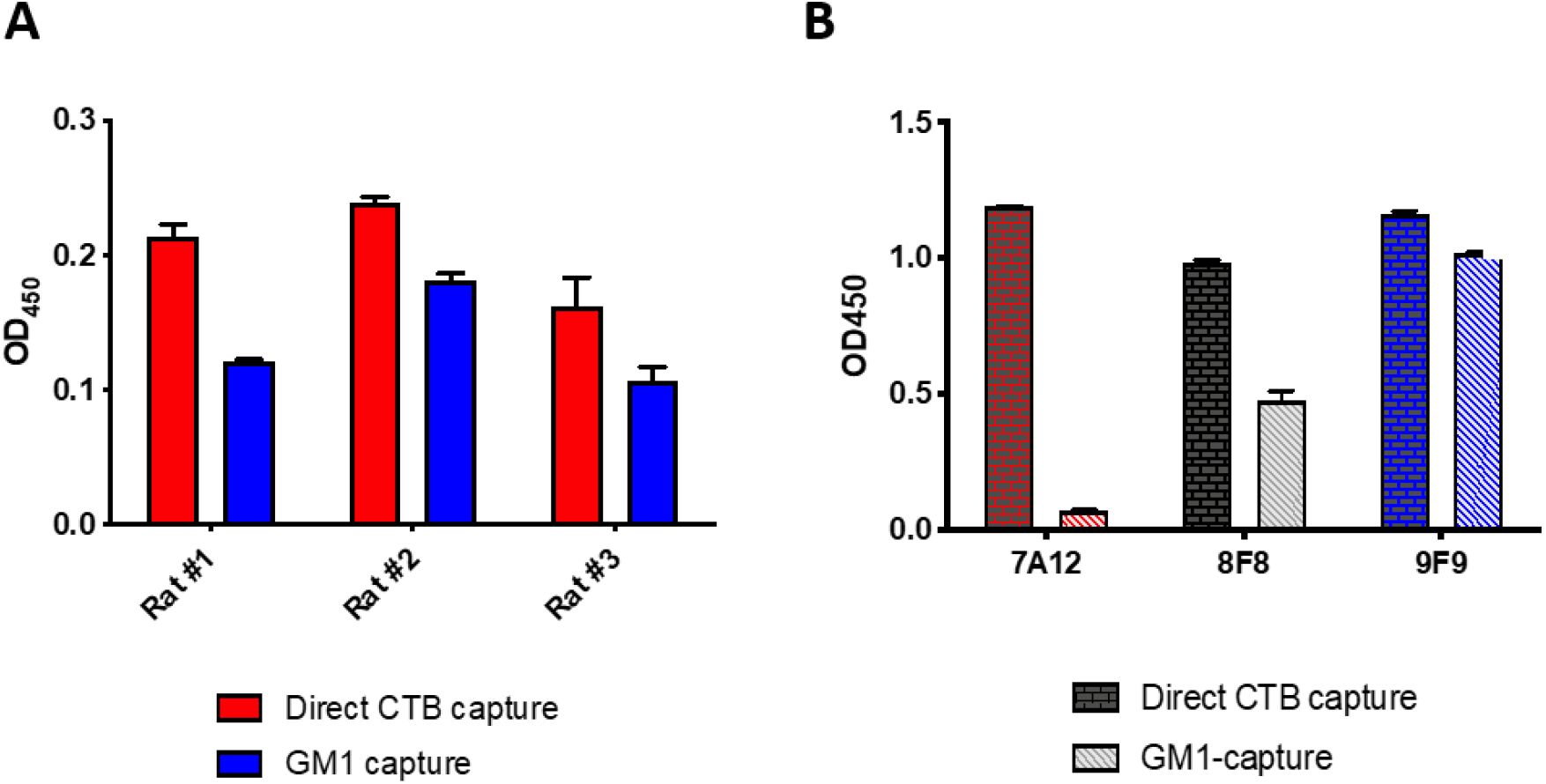
Generation and screening of hybridoma supernatants reacting to CTB. **A.** Three Wistar rats were immunized with gCTB to generate a panel of mAbs and blood serum from each rat was diluted and analyzed for the level of anti-CTB IgG antibodies by direct CTB-capture and GM1-capture ELISA. **B.** Characterization of three hybridomas, 7A12, 8F8 and 9F9, which showed high anti-CTB antibody levels by ELISA. The hybridoma cell culture supernatants showed distinctive binding patterns to CTB in direct CTB-capture vs. GM1-capture ELISA. All three hybridoma showed similarly high binding in CTB-capture ELISA. By contrast, 7A12 showed the least and negligible binding while 9F9 showed the highest binding in GM1-capture ELISA. Bars represent means ± SD (N=2).

### The mAbs 7A12B3 and 9F9C7 bind CTX, CTB, and EPICERTIN with high affinities

To determine the antigen-binding properties of 7A12B3 and 9F9C7 mAbs, we initially performed ELISA wherein CTB, CTX, and LTB were directly coated on the plates. Both 7A12B3 and 9F9C7 mAbs showed similar high binding affinities to CTB, expressed as half-maximal effective concentrations (EC_50_) of 0.028 ± 0.003 and 0.036 ± 0.003 μg/mL, respectively (**Figure 2A**). Likewise, 7A12B3 and 9F9C7 mAbs bound to CTX with high binding affinities represented by low EC_50_ values (0.017 ± 0.003 and 0.041 ± 0.005 μg/mL, respectively). The 9F9C7 mAb showed slightly lower affinity to CTX than 7A12B3, possibly due to marginal occlusion of the epitope by the holotoxin A subunit (**Figure 2B**). Despite LTB’s high amino acid sequence similarity to CTB,^34,35^ both 7A12B3 and 9F9C7 mAbs showed a substantially lower affinity to LTB than to CTB and CTX, with an EC_50_ value of 1.109 ± 0.162 and 135 ± 70 μg/mL, respectively (**Figure 2C**). Average EC_50_ values of 7A12B3 and 9F9C7 mAbs binding to all three molecules are presented in **Table 1**.

**Figure 2.**
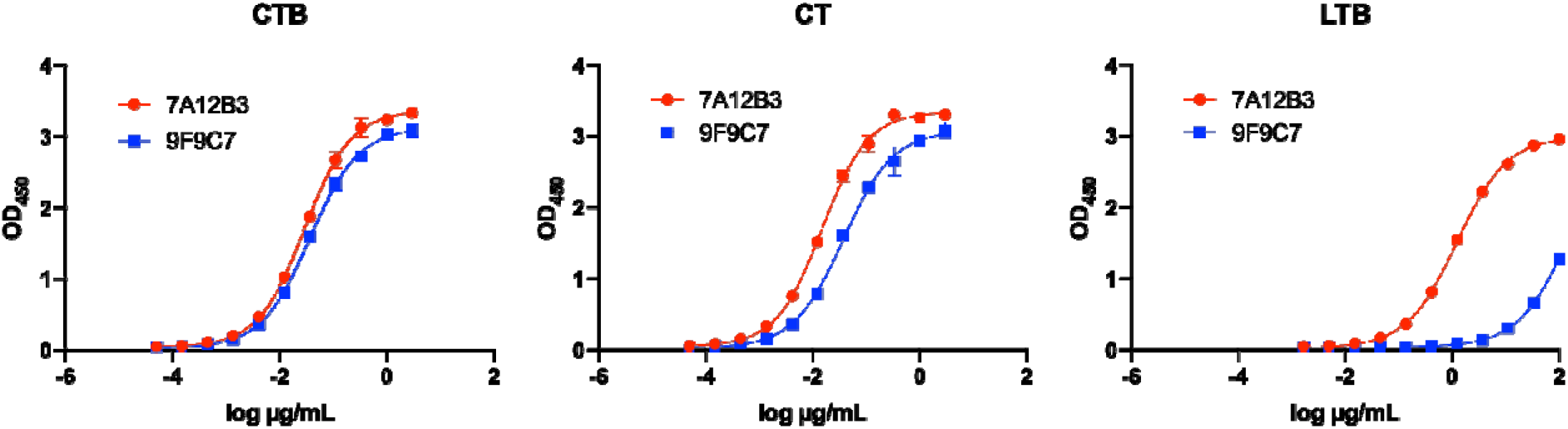
Analysis of 7A12B3 and 9F9C7 mAbs binding to CTX, CTB and LTB in antigen-capture ELISA. ELISA plates were coated with 2 μg/mL of CTB, CTX or the *Escherichia coli* heat-labile enterotoxin B subunit (LTB). Three-fold serially diluted 7A12B3 or 9F9C7 mAbs (3,000 – 0.051 ng/mL for CTB and CTX; 100,000 – 1.69 ng/mL for LTB) were added to the plates and incubated, and plate-bound mAbs were detected with an anti-rat IgG secondary antibody. Representative graphs are shown. The assays were performed in triplicate, and each data point represents the mean ± SD. Data were analyzed and plotted using the GraphPad Prism 8 software and obtained from at least two independent experiments. The half-maximal effective concentrations (EC_50_s) were determined by nonlinear regression analysis (GraphPad Prism 8) and displayed in Table 1.

**Table 1.** Average EC_50_ values and standard deviation (N=2) of 7A12B3 and 9F9C7 mAbs binding to CTX, CTB and LTB

| EC<sub>50</sub> (µg/mL) |  |  |  |
| --- | --- | --- | --- |
|  | CTX | CTB | LTB |
| **7A12B3 mAb** | 0.017 ± 0.003 | 0.028 ± 0.003 | 1.09 ± 0.162 |
| **9F9C7 mAb** | 0.041 ± 0.005 | 0.036 ± 0.003 | >100 |

To further dissect the antigen-binding profiles of 7A12B3 and 9F9C7, SPR analysis was employed, in which each mAbs was immobilized on a CM5 sensor chip while CTX, CTB, EPICERTIN, and LTB were used as soluble analytes. **Figure 3** shows representative sensorgrams. Analysis of binding kinetics revealed that 7A12B3 mAb had an average association rate constant (*k*_on_) of 2.1 x 10^6^ (1/Ms), a dissociation rate constant (*k*_o_ff) of 1.8 x 10^-4^ (1/s), and an average equilibrium dissociation constant (*K*_D_) of 83.0 pM to CTX. This mAb also showed similar high binding affinity to CTB and EPICERTIN, with average *K*_D_ values of 88.9 and 158.8 pM, respectively (**Figure 3A)**. On the other hand, 9F9C7 mAb showed slower association and slower dissociation for CTX and CTB compared to 7A12B3. The 9F9C7 mAb also showed slower association but similar dissociation to EPICERTIN, compared to 7A12B3 (**Figure 3B)**. Thus, 9F9C7 mAbs turned out to have overall lower binding affinities to the three analytes, as 9F9C7 showed an average *K*_D_ of 3.9 nM to CTX, 4.8 nM to CTB, and 32.4 nM to EPICERTIN. These values correspond to approximately 50 times (CTX and CTB) and 200 times (EPICERTIN) lower affinity when compared to 7A12B3. In sharp contrast, neither mAbs showed measurable binding to LTB under the conditions used in this SPR analysis (**Figure 3A and B)**. Average association and dissociation rate constants and *K*_D_ values are summarized in **Table 2**.

**Figure 3.**
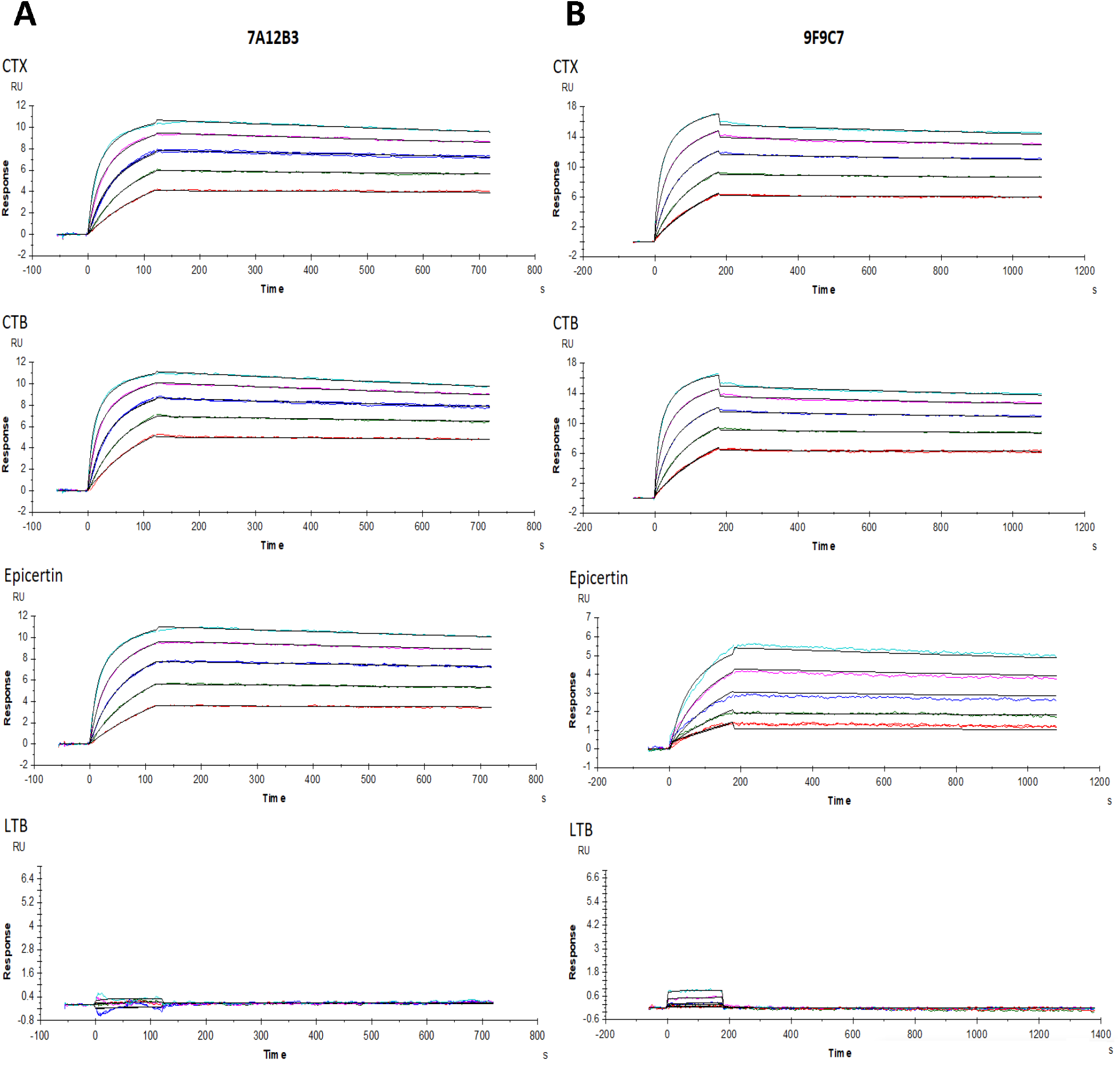
SPR analysis of 7A12B3 and 9F9C7 mAbs binding affinities to CTX, CTB, EPICERTIN and LTB. Each mAb was immobilized on a CM5 sensor chip. Each analyte (CTX, CTB, EPICERTIN, and LTB) was tested in a range of concentrations (0, 1.68, 3.37, 6.73, 13.47, 26.9375 nM) against immobilized 7A12B3 mAb or 9F9C7 mAb (0, 53.1, 106.3, 212.5, 425, 850, 1700 nM). Representative sensorgrams for (A) 7A12B3 and (B) 9F9C7 are shown. Each experiment was conducted with at least three replicates.

**Table 2.** Association and dissociation constants of CT, CTB and EPICERTIN binding to 7A12B3 and 9F9C7 mAbs generated by surface plasmon resonance.

| 7A12B3 mAb |  |  |  |  |
| --- | --- | --- | --- | --- |
| | $k_{on}$ (1/Ms) | $k_{off}$ (1/s) | $K_D$ (pM) | Chi<sup>2</sup>(RU<sup>2</sup>) |
| **CTX** | $2.1 \times 10^6$ ($2.0 \times 10^6 - 2.4 \times 10^6$) | $1.8 \times 10^{-4}$ ($1.5 \times 10^{-4} - 2.1 \times 10^{-4}$) | 83.0 (72.9 – 100.0) | $7.6 \times 10^{-3}$ ($5.8 \times 10^{-3} - 8.8 \times 10^{-3}$) |
| **CTB** | $2.6 \times 10^6$ ($2.4 \times 10^6 - 2.8 \times 10^6$) | $2.3 \times 10^{-4}$ ($2.0 \times 10^{-4} - 2.6 \times 10^{-4}$) | 88.9 (81.1 – 103.0) | $10.0 \times 10^{-3}$ ($7.2 \times 10^{-3} - 16.0 \times 10^{-3}$) |
| **EPICERTIN** | $1.1 \times 10^6$ ($1.0 \times 10^6 - 1.2 \times 10^6$) | $1.7 \times 10^{-4}$ ($1.6 \times 10^{-4} - 1.7 \times 10^{-4}$) | 158.0 (145.0 – 171.0) | $6.0 \times 10^{-3}$ ($4.1 \times 10^{-3} - 8.0 \times 10^{-3}$) |

| 9F9C7 mAb |  |  |  |  |
| --- | --- | --- | --- | --- |
| | $k_{on}$ (1/Ms) | $k_{off}$ (1/s) | $K_D$ (nM) | Chi<sup>2</sup>(RU<sup>2</sup>) |
| **CTX** | $2.3 \times 10^4$ ($2.1 \times 10^4 - 2.6 \times 10^4$) | $9.1 \times 10^{-5}$ ($8.4 \times 10^{-5} - 10.0 \times 10^{-5}$) | 3.9 (3.7 – 4.1) | $8.3 \times 10^{-3}$ ($7.3 \times 10^{-3} - 9.3 \times 10^{-3}$) |
| **CTB** | $2.0 \times 10^4$ ($1.9 \times 10^4 - 2.2 \times 10^4$) | $9.8 \times 10^{-5}$ ($9.4 \times 10^{-5} - 10.0 \times 10^{-5}$) | 4.8 (4.6 – 5.2) | $9.9 \times 10^{-3}$ ($7.4 \times 10^{-3} - 11.6 \times 10^{-3}$) |
| **EPICERTIN** | $4.9 \times 10^3$ ($4.9 \times 10^3 - 5.0 \times 10^3$) | $1.6 \times 10^{-4}$ ($1.5 \times 10^{-4} - 1.8 \times 10^{-4}$) | 32.4 (30.0 – 36.3) | $3.0 \times 10^{-2}$ ($2.8 \times 10^{-2} - 3.1 \times 10^{-2}$) |

### The 7A12B3 mAb, but not 9F9C7, blocks CTB binding to its receptor GM1

A GM1-capture ELISA was initially conducted to analyze the impact of 7A12B3 and 9F9C7 mAbs on the receptor binding activity of CTB. Consistent with our observations from the culture supernatants of the parental hybridoma clones (**Figure 1B)**, the 7A12B3 mAb markedly blocked the binding of CTB to GM1 whereas no significant effect was observed in the presence of 9F9C7 mAb (data not shown). To analyze more rigorously the blocking properties of 7A12B3 mAb in CTB-GM1 interaction, we used a GM1-capture, KDEL-detection (GM1/KDEL) ELISA, in which EPICERTIN bound to the glycosphingolipid receptor was detected using anti-KDEL mAb.^22^ The 7A12B3 mAb at 0.1 – 1 μg/mL dose-dependently inhibited the binding of EPICERTIN to GM1 (**Figure 4A, left panel**). In contrast, 9F9C7 mAb showed a relatively smaller effect on EPICERTIN binding to GM1 and only at a concentration of 1 μg/mL shifted the EPICERTIN binding curve to a level very similar to that observed with 0.1 μg/mL of 7A12B3 mAb (**Figure 4A, right panel**). In a competitive GM1/KDEL ELISA in which varying concentrations of respective mAbs were pre-incubated with a fixed concentration of EPICERTIN at 100 ng/mL, the IC_50_ values for 7A12B3 and 9F9C7 mAbs on the EPICERTIN binding to GM1 were determined to be 201.2 vs. 993.7 ng/mL respectively (**Figure 4B**).

**Figure 4.**
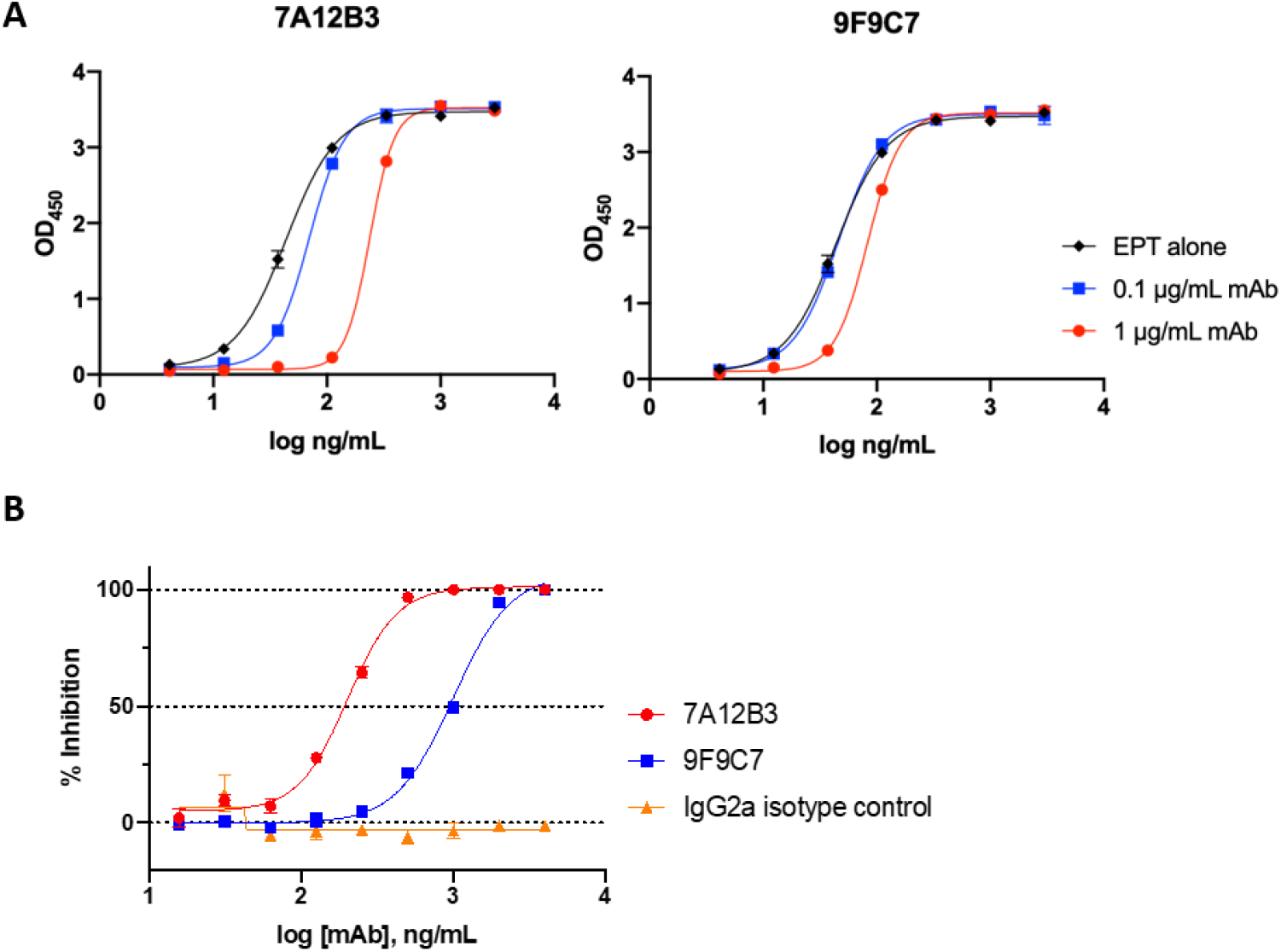
The 7A12B3 mAb, but not 9F9C7, blocks CTB binding to GM1 ganglioside. **A.** Impacts of 7A12B3 and 9F9C7 mAbs on the EPICERTIN binding curve in GM1/KDEL ELISA. The 7A12B3 mAb concentration-dependently blocked the interaction of CTB with GM_1_ at 0.1 – 1 μg/mL, whereas the 9F9C7 mAb had much less effects. **B**. A competitive ELISA was employed to determine the effect of mAbs on EPICERTIN - GM1 interaction. Varying concentrations of respective mAbs, including a rat IgG2a isotype control, were pre-incubated with 100 ng/mL of EPICERTIN and applied to ELISA plates coated with GM1. Plate-bound EPICERTIN was detected with an anti-KDEL mAb, as described previously. Half maximal inhibitory concentration (IC50) of 7A12B3 and 9F9C7 mAbs were determined by a non-linear regression analysis using GrapPad Prism 8. Data were obtained from at least three independent experiments, and representative graph from one experiment is shown. Each data point represents the mean ± SD of duplicate samples.

### The 7A12B3 mAb effectively inhibits CTX-induced cAMP in Caco2 cells

The inhibitory effects of 7A12B3 and 9F9C7 mAbs on the biological functions of CTX were analyzed in the Caco2 cell line model of cAMP induced by CTX. **Figure 5A** shows that CTX (0.5 μg/mL) preincubated with 7A12B3 mAb (1 μg/mL) induced a significantly lower level of cytoplasmic cAMP in Caco2 cells when compared to CTX alone (86.7% inhibition; *p* < 0.0001), whereas 9F9C7 mAb showed significant yet less inhibitory effect (62.6% inhibition; *p* = 0.0032). The inhibitory effect of 7A12B3 mAbs was significantly different from that of 9F9C7 mAb *(p* = 0.0274) or a rat IgG2a isotype control *(p* = 0.0002). The marginal inhibition observed with a rat IgG2a isotype control antibody was not statistically significant from the PBS vehicle control (*p*=0.0626). The inhibitory effects of 7A12B3 and 9F9C7 mAbs on CTX-induced elevation of cAMP in Caco2 cells were concentration dependent (**Figure 5B**). When CTX was co-incubated with 1 μg/mL of mAbs, 7A12B3 inhibited the induction of cAMP by 88.1%, whereas significantly less inhibition was observed with 9F9C7 mAb (66.8%). Both mAbs had minimal inhibitory effects at 0.25 μg/mL (14% vs. 12%, respectively), which were indistinguishable from the background effects of a Rat IgG isotype control antibody.

**Figure 5.**
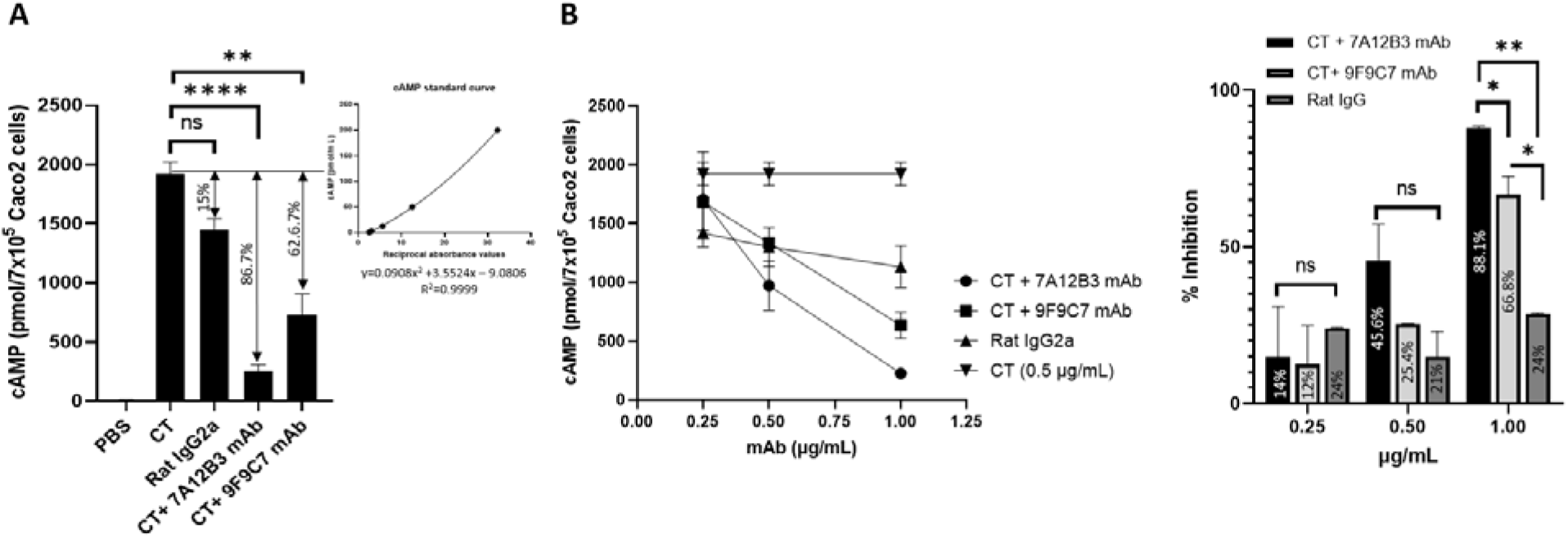
The 7A12B3 mAb effectively inhibits CTX-induced cAMP levels in Caco2 cells. **A.** The inhibitory effects of 7A12B3 and 9F9C7 mAbs (0.25-1.0 μg/mL) on CTX-induced cAMP were evaluated under preincubation of the mAbs with CTX (0.5 μg/mL) for 20 minutes. The 7A12B3 mAb strongly inhibited CTX-induced cAMP levels in Caco2 cells, whereas 9F9C7 had reduced inhibitory effects. \*\*\*\**p* < 0.0001, \*\**p* < 0.01, one-way measures ANOVA with Bonferroni’s multiple comparisons tests. Inset shows a standard curve of cAMP, with a non-linear regression analysis (GrapPad Prism 8) used to determine cAMP values in samples. **B**. Concentration-dependent inhibition of CTX-induced cAMP levels in Caco2 cells by 7A12B3 and 9F9C9 mAbs. \**p* < 0.05, \*\**p* < 0.01, two-way measures ANOVA with Bonferroni’s multiple comparisons tests. Data obtained from two independent experiments. Each data point represents the mean ± SD of triplicate samples.

### The 9F9C7 mAb effectively detects CTB docking on the surface of target cells

Flow cytometric analysis was conducted to evaluate the utility of 7A12B3 and 9F9C7 mAbs to detect CTB and its variants bound to the surface of target cells. Although 7A12B3 was able to detect EPICERTIN on the surface of Caco2 epithelial cells, 9F9C7 showed superior detectability with increased fluorescence signal at the same concentration used (**Figure 6A, left panel**). Thus, the latter mAb was used in further analysis.

**Figure 6.**
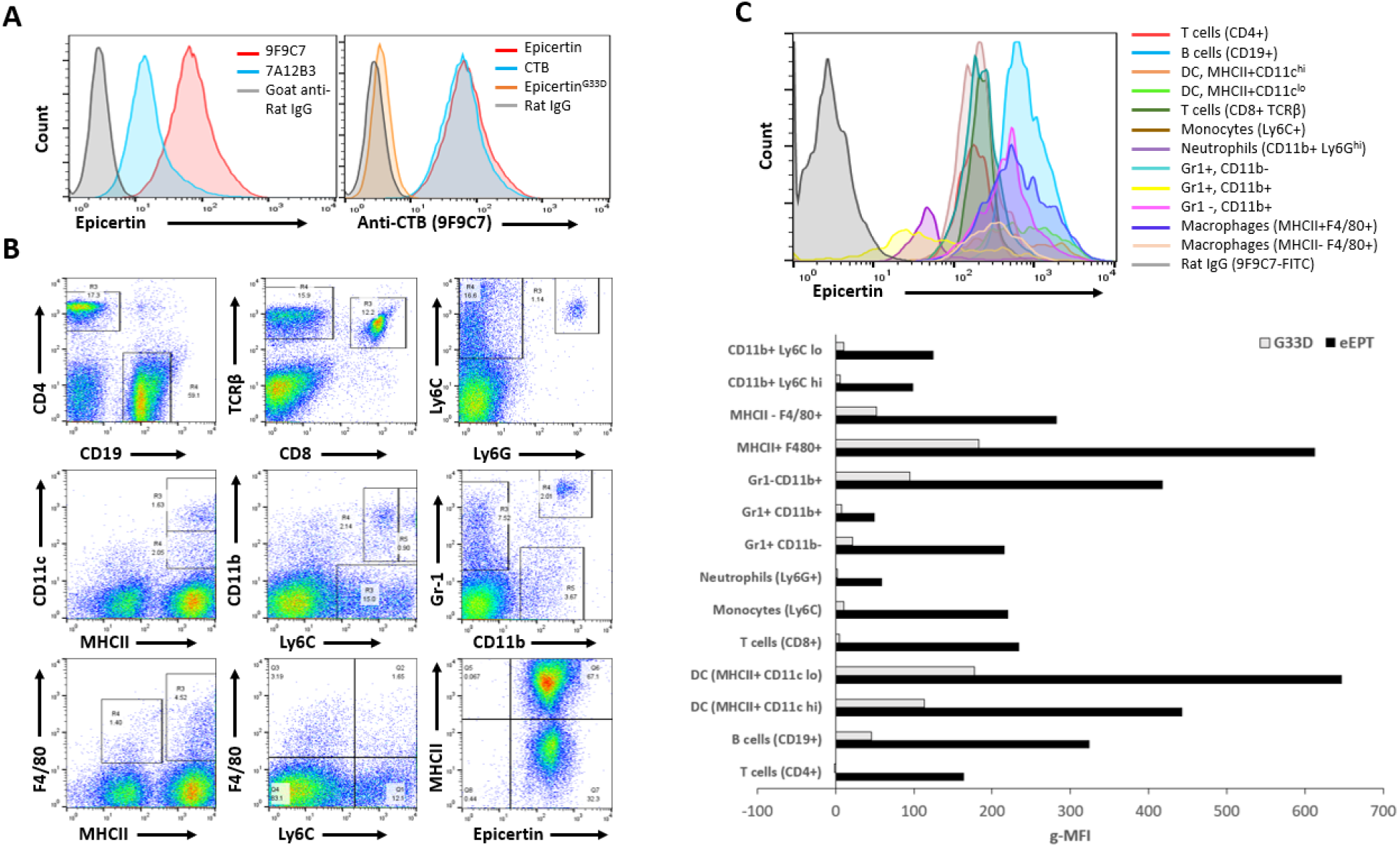
The 9F9C7 mAb detects EPICERTIN but not EPICERTIN^G33D^ bound to the surface of human Caco2 cell line and to mouse spleen leukocytes. **A.** Flow cytometry analysis of EPICERTIN binding to Caco2 cells detected with 2 μg/mL of 7A12B3 or 9F9C7 mAbs followed by FITC-labeled goat-anti-rat IgG (left panel). Detection of EPICERTIN, CTB and EPICERTIN^G33D^ (2 μg/mL) binding to human Caco2 cell line detected with anti-CTB 9F9C9 mAb (right panel). **B.** Flow cytometric analysis of EPICERTIN and EPICERTIN^G33D^ binding to spleen leukocytes stained with a panel of fluorochrome-labeled antibodies and FITC-conjugated 9F9C7 mAb. **C**. Histograms and geometric mean fluorescence intensity of EPICERTIN bound to the mouse spleen leukocytes gated in panel B.

Using the 9F9C7 mAb, we observed strong and comparable binding of EPICERTIN and native CTB (nCTB) to the surface of Caco2 cells, whereas EPICERTIN^G33D^, a variant which lacks GM1-binding activity, was only marginally detected on the cell surface (**Figure 6A, right panel**). Next, we attempted to characterize EPICERTIN’s target immune cells using 9F9C7. To this end, mouse spleen cells were incubated with EPICERTIN or EPICERTIN^G33D^, stained with different combination of cell-surface marker-specific antibodies and 9F9C7, and gated to sort target cell subpopulations, as shown in **Figure 6B**. The geometric mean fluorescence intensity (G-MFI) of EPICERTIN and EPICERTIN^G33D^ detected with FITC-labeled goat anti-Rat IgG antibody above the background levels generated by this second antibody alone is shown in **Figure 6C.** We found that EPICERTIN efficiently bound to the surface of all myeloid and lymphoid cells analyzed, with major histocompatibility complex class II (MHC II)-positive dendritic cells and macrophages being the most prominent targets. Surprisingly, EPICRTIN^G33D^ appeared to recognize some of the cell types, including B cells, monocytes, macrophages, and dendritic cells, although the degrees of binding to these cells were overall much lower compared to EPICERTIN (**Figure 6C**).

### The 9F9C7 mAb detects EPICERTIN by immunohistochemistry on frozen colon sections

Fluorescent immunohistochemistry was conducted using FITC-conjugated 9F9C7 mAb to detect and locate EPICERTIN bound to the surface of colon epithelial cells upon oral administration of the protein in mice. As shown in **Figure 7**, EPICERTIN was detected on frozen colon tissue sections from 6 to 24 hr after oral administration. Of note, the fluorescence signal was consistently detected on epithelial cells within the colonic glands while not prominent on epithelial cells facing the luminal side of the colon (**Figure 7**). The staining of EPICERTIN delineated the luminal side of differentiated crypt epithelial cells (near the colon crypt opening) and less differentiated epithelial cells located at the bottom of the crypts. Additionally, the image disclosed that EPICERTIN’s fluorescence signal at the plasma membrane seem to follow closely that of the adhesion molecule E-cadherin (although not overlapping) in less differentiated and mucin-rich crypt epithelial cells. Prominent EPICERTIN staining was also observed in the cytoplasm of those cells. Meanwhile, FITC-9F9C7 mAb did not show any fluorescence signal in colon tissues from EPICERTIN-untreated mice, confirming the specificity of the antibody.

**Figure 7.**
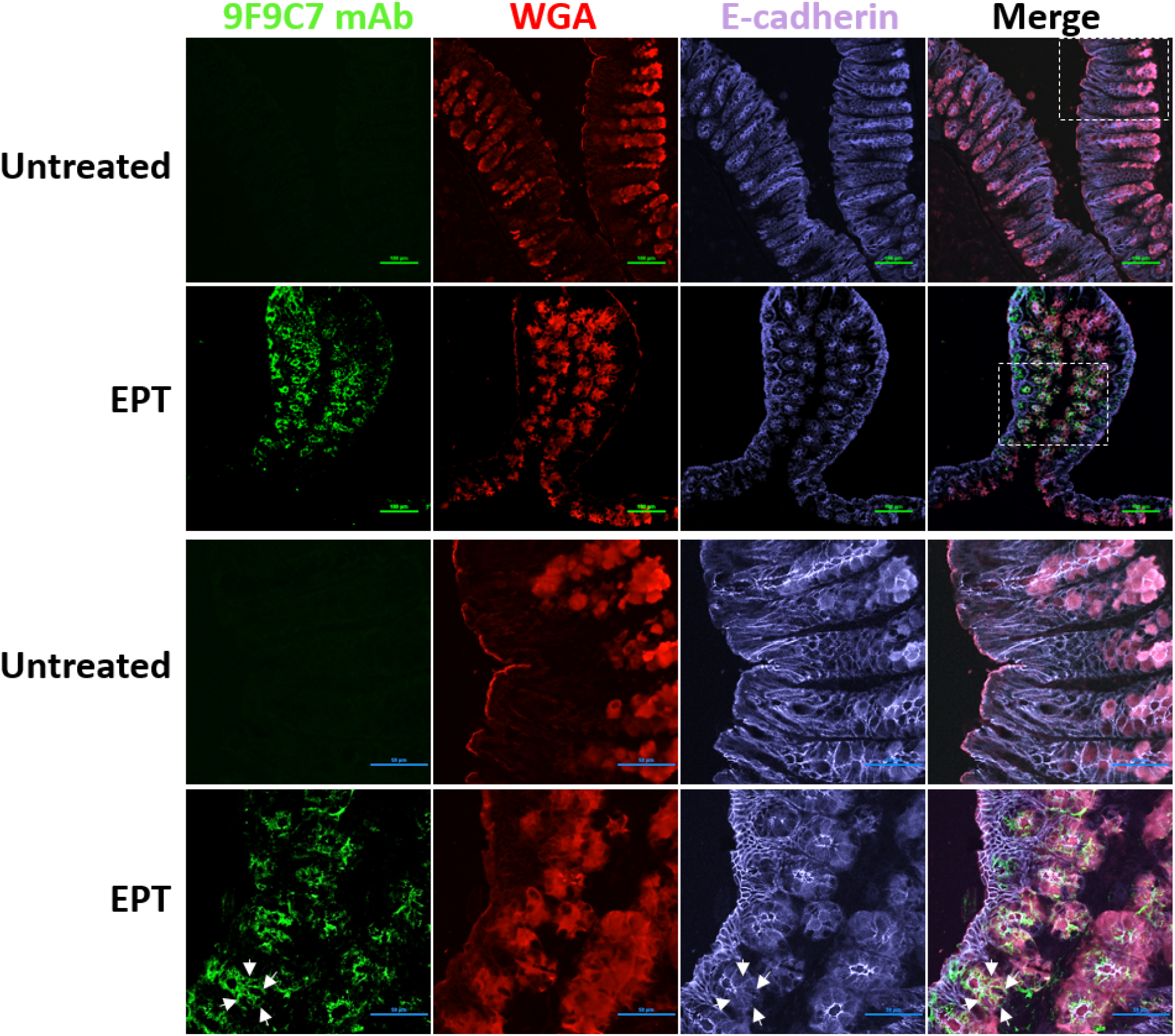
IHC detection of EPICERTIN bound to the surface epithelial cells of mouse colon tissue. Confocal laser scanning microcopy of colon tissues from mice treated or left untreated with EPICERTIN by oral gavage. Colon tissues were collected at different time pointes after oral administered Epicertin and embedded in OCT compound for cryosectioning. Seven microns tick tissue sections were made and stained with 5 μg/mL Pheno Vue Fluor 594-WGA, 2 μg/mL anti-E-cadherin and 2 μg/mL anti-CTB (FITC-9F9C7 mAb). White arrows indicate distribution of EPT at the plasma membrane of individual colon crypt epithelial cells. Images were collected with a Nikon A1R Confocal laser scanning microscope using 20 × (top panel) and 60 × (lower panel) magnification lenses with appropriate channels and the data processed with the NIS Elements imaging software. Areas delineated by dotted line squares in panel A correspond to high magnification images shown in lower panel.

## DISCUSSION

Since the introduction of hybridoma technology over 45 years ago,^30^ several anti-CTX mAbs targeting different epitopes on the A and B subunits have been generated in early studies.^36–41^ Some of those mAbs recognized the GM1 receptor binding site of CTB or showed distinctive neutralizing CTX activity,^42^ whereas other mAbs that were generated against CTX peptides, often resulted in the generation of mAbs with polyspecific binding properties or completely lacked CTX binding activity.^43,44^ They aided in building the current understanding of CTX secretion, assembly^45,46^, endocytosis and intoxication,^47^ and were also instrumental in understanding the potent immunogenicity of CTB and the structurally homologous LTB.^37,48^ However, most of those anti-CTB mAbs were characterized only using outdated immunoassay-based methods, providing limited information about their binding profiles. Recently, novel recombinant CTB variant and fusion molecules have been generated, some of which were found to have unique biological functions, such as mucosal healing promoted by a CTB variant containing an ER retention motif, EPICERTIN.^21,33^ To aid in the preclinical development of CTB-based vaccines and biotherapeutics, we attempted to isolate and characterize new anti-CTB mAbs that are suitable for mechanistic investigations and pharmacological studies.

The 7A12 and 9F9 hybridoma cell culture supernatants were found to bind CTB with distinctive features in both direct CTB-capture and GM1-capture ELISAs (**Figure 1**). Both mAbs bound CTB with high affinity, but only 9F9 effectively bound CTB in GM1-capture ELISA, indicating that 7A12 recognizes an epitope near or within the region of CTB responsible for GM1 interaction. On the other hand, the 9F9 hybridoma supernatant appeared to recognize a distinct epitope most probably not involved in GM1 binding. These results demonstrate that our screening procedure employed here successfully led to the isolation of two mAbs with distinct CTB-binding profiles in terms of reactivity with the antigen’s GM1-receptor binding site.

The 7A12B3 and 9F9C7 mAbs were found to bind the native CTX and CTB with similar binding affinities in direct ELISA (**Figure 2**, **Table 1**). However, SPR analysis revealed that these mAbs have distinct binding kinetics. The overall binding affinity of 7A12B3 mAb was higher than that of 9F9C7 mAb; 158.0 pM vs. 32.4 nM for EPICERTIN, 83.0 pM vs. 3.9 nM for CTX, and 88.9 pM vs. 4.8 nM for CTB, respectively (**Figure 3**, **Table 2**). In contrast, neither 7A12B3 or 9F9C7 bound to LTB in SPR (**Figure 3**), along the lines of the ELISA data that also showed substantially low affinity of these mAbs to LTB compared to CTB and CTX (**Figure 2**). These results demonstrate the exquisite specificity of 7A12B3 and 9F9C7 mAbs to CTB, given that LTB has high (~84%) amino-acid sequence homology with CTB.^34,35^ The binding affinity of 7A12B3 to EPICERTIN was slightly lower than to CTX or CTB, and those differences might be explained by the Asn4→Ser mutation and/or the presence of C-terminal extension comprised of the hexapeptide SEKDEL sequence in EPICERTIN. Of note, 7A12B3 mAb but not 9F9C7 effectively inhibited the binding of EPICERTIN to GM1 ganglioside (**Figure 4**), strengthening the idea that the former recognizes an epitope near the GM1 binding site of CTB, whereas the latter is relatively indifferent to CTB-GM1 interaction.

CTX induces cAMP overproduction in the cytoplasm of target cells. Our data demonstrated that the 7A12B3 mAb has strong CTX-neutralizing effects, almost completely inhibiting the cytoplasmic accumulation of cAMP induced by CTX in Caco2 cells (**Figure 5B**). Interestingly, even though 9F9C7 mAb appeared to bind to an epitope distal to the GM1-binding site, we found that the mAb was also able to inhibit the effects of CTX on the elevation of cytoplasmic cAMP in Caco2 cells, although at lower levels than 7A12B3 mAb. We speculate that 9F9C7 mAb may form complexes with CTX in solution, which in turn collaterally compromises CTX-GM1 interaction and/or entry to target cells.

Based on the results from the competitive ELISA (**Figure 4**) and CTX cAMP reporter assays (**Figure 5**), 7A12B3 was thought to target an epitope proximal to the GM1 binding site, an area of CTB that would be occluded after engaging the cell-surface glycosphingolipid receptor. However, flow cytometry analysis revealed that the mAb is capable of detecting EPICERTIN on the surface of Caco2 epithelial cells (**Figure 6A, left panel**). Nevertheless, 9F9C7, which was selected based on effective recognition of GM1-bound CTB (**Figure 1B**), showed superior detectability of cell-bound EPICERTIN and thus justified the use of this mAb to explore the target cell binding profile of EPICERTIN. The flow cytometry analysis (**Figure 6**) revealed that EPICERTIN and CTB equally bound to the surface of Caco2 cells, as anticipated from their similar binding affinity to GM1 ganglioside.^19^ In sharp contrast, EPICERTIN^G33D^ was only marginally detected on the cell surface (**Figure 6A, right panel**), suggesting that the glycosphingolipid is the primary receptor for EPICERTIN in the colon epithelial cell line. To our surprise, however, we found inconsistent binding patterns of EPICERTIN and EPICERTIN^G33D^ in mouse spleen leukocytes. For instance, EPICERTIN’s geometric mean fluorescence intensity (gMFI) ranged between 50-300 whereas the gMFI of EPICERTIN^G33D^ ranged from 0-45 (**Figure 6B, C**). Although GM1 ganglioside has been long considered the sole receptor for CTB binding and internalization by epithelial cells, recent findings pointed to the presence of alternative receptors, such as fucosylated glycoconjugates.^49–51^ In addition, cycling of KDEL receptors between the Golgi and cell membrane^52^ could partly account for the cellular binding patterns of EPICERTIN and the G33D variant. Thus, differential expression of those receptors might explain CTB binding to leukocytes in a cell type specific manner. Nevertheless, because the degree of binding was overall substantially higher with EPICERTIN than with the non-GM1-binding counterpart, it seems reasonable to assume that EPICERTIN’s effects on immune cells are likely mediated by GM1 receptor engagement.

The expression of GM1 is not limited to intestinal epithelial cells. It is expressed in a variety of other cell types, including cortical and peripheral neurons^53,54^ and leukocytes,^55^ among others. A differential expression of GM1 on human monocytes suggested the presence of two monocytes subpopulations with functional differences in terms of endocytic activity and lipopolysaccharide responsiveness in peripheral blood.^56^ CTB is known to bind to GM1 expressed on the surface of leukocytes, particularly innate immune cells such as dendritic cells, macrophages and B cells, which are the major antigen-presenting cells.^8^ CTB binding to GM1 on B cells was associated with cAMP-independent inhibition of mitogen-stimulated B cell proliferation and enhanced expression of MHCII molecules,^26,57^ whereas binding of CTB on T lymphocytes was found to inhibit mitogen or antigen-induced T-cell proliferation.^26^ Of note, however, the nature of the enhanced immune responses to antigens coupled to CTB and the dampening of autoimmune responses by this protein are still largely unknown. In the case of antibody-mediated immune responses against infectious microorganisms, the increased MHC II expression on B cells induced by CTB might partially explain the immunomodulatory effect favoring this outcome.^8^ In the case of suppression of airway allergic inflammation, CTB’s therapeutic effect appeared to reside in its capacity to reprogram dendritic cells to instruct B cells for IgA class switch.^58^ As shown in **Figure 6**, EPICERTIN highly bound to antigen-presenting cells compared to other leukocytes, particularly MHC II^+^ CD11c^lo^ dendritic cells, MHC II^+^ F480^+^ macrophages and CD19^+^ B cells. Although it remains a matter of speculation at this point, such preferential binding may suggest EPICERTIN’s distinctive effects on these cells that could have implications for the protein’s immunomodulatory effects.

The specific interaction of CTB with GM1 ganglioside expressed on the surface of intestinal epithelial cells is a well-known mechanism responsible for the internalization of CTX and its virulence during *V. cholera* infection.^59^ This high affinity interaction has been exploited in vaccine development where CTB is used as an adjuvant and carrier protein. Additionally, the ability of CTB to undergo retrograde transportation in target cells may provide opportunities for the development of novel pharmaceutical products with unique biological functions, as exemplified by EPICERTIN, which was found to be retained in the ER of colon epithelial cells where it induces an unfolded protein response leading to epithelial repair activity. However, the type of colon epithelial cell targeted/responsible for such a response remaines to be identified. In the IHC analysis on cryosections of mouse colon tissue using the 9F9C7 mAb (**Figure 7**), we were able to clearly detect EPICERTIN in the colon at 6 hr and up to 24 hr after oral administration. Interestingly, EPICERTIN was detected mainly on the surface of epithelial cells lining the openings of colonic crypts with consistent detection on less differentiated cells at the bottom of the crypts, including crypt-resident goblet cells that are densely stained with the WGA lectin^60^. This observation suggests that EPICERTIN might have prominent effects on the colon stem cell compartment with proliferative capability than on differentiated epithelial cells. However, this conjecture needs further verification as we cannot rule out the possibility that the detection of EPICERTIN mostly in the crypt base region might be a procedural artifact during the flushing procedure of colons before tissue embedding, which could have inadvertently removed EPICERTIN bound to the inter-crypt epithelium exposed on the luminal side. Our future study will address this issue by further IHC analysis of ex vivo-cultured mouse and human colon tissues.

In conclusion, two novel mAbs were generated that bind CTX, CTB and EPICERTIN with high affinity and specificity. The 7A12B3 mAb effectively inhibited the binding of CTB to GM1 and neutralized CTX, whereas the 9F9C7 mAb showed superior capacity to detect EPICERTIN binding to the surface of target cells. Coupled with our earlier reports showing the utility of 9F9C7 in immunofluorescence and confocal microscopy^23^ and 7A12B3 in rodent pharmacokinetic analysis of EPICERTIN^25^, these mAbs provide valuable tools to facilitate the investigation and development of CTB variants as novel biopharmaceutical candidates.

## Acknowledgments

We thank Wendy Cecil and Micaela Reeves for critical reading of the manuscript. The work was supported by a grant from the Leona M. and Harry B. Helmsley Charitable Trust (2014PG-MED001) and a U.S. National Institutes of Health grant (R01 DK123712) to NM.

## Data availability

The datasets generated during the current study are available from the corresponding author on reasonable request.

## Author contributions statement

NM and NVG designed the study, conducted experiments, carried out data analysis, data interpretation, and drafted the manuscript. ICSC and MD conducted experiments and helped to draft the manuscript.

## References

1 Harris, J., LaRocque, R., Qadri, F., Ryan, E. & Calderwood, S. Cholera. Lancet 379, 2466–2476, doi:10.1016/S0140-6736(12)60436-X (2012).

2 Cuatrecasas, P. Interaction of Vibrio cholerae enterotoxin with cell membranes. Biochemistry 12, 3547–3558 (1973).

3 Holmgren, J., Lönnroth, I. & Svennerholm, L. Fixation and inactivation of cholera toxin by GM1 ganglioside. Scandinavian journal of infectious diseases 5, 77–78 (1973).

4 Merritt, E.A. et al. Crystal structure of cholera toxin B pentamer bound to receptor GM1 pentasaccharide. Protein Science 3, 166–175 (1994).

5 Holmgren, J., Czerkinsky, C., Lycke, N. & Svennerholm, A.-M. Strategies for the induction of immune responses at mucosal surfaces making use of cholera toxin B subunit as immunogen, carrier, and adjuvant. The American journal of tropical medicine and hygiene 50, 42–54 (1994).

6 Azegami, T., Itoh, H., Kiyono, H. & Yuki, Y. Novel transgenic rice-based vaccines. Archivum immunologiae et therapiae experimentalis 63, 87–99 (2015).

7 Baldauf, K.J., Royal, J.M., Hamorsky, K.T. & Matoba, N. Cholera toxin B: one subunit with many pharmaceutical applications. Toxins 7, 974–996 (2015).

8 Stratmann, T. Cholera Toxin Subunit B as Adjuvant--An Accelerator in Protective Immunity and a Break in Autoimmunity. Vaccines (Basel) 3, 579–596, doi:10.3390/vaccines3030579 (2015).

9 Solbreux, P.M., Dive, C. & Vaerman, J.-P. Anti-cholera toxin IgA-, IgG-and IgM-secreting cells in various rat lymphoid tissues after repeated intestinal or parenteral immunizations. Immunological investigations 19, 435–451 (1990).

10 Apter, F., Lencer, W., Finkelstein, R., Mekalanos, J. & Neutra, M. Monoclonal immunoglobulin A antibodies directed against cholera toxin prevent the toxin-induced chloride secretory response and block toxin binding to intestinal epithelial cells in vitro. Infection and immunity 61, 5271–5278 (1993).

11 Sanchez, J., Johansson, S., Löwenadler, B., Svennerholm, A. & Holmgren, J. Recombinant cholera toxin B subunit and gene fusion proteins for oral vaccination. Research in microbiology 141, 971–979 (1990).

12 Vendetti, S. et al. Polyclonal Treg cells enhance the activity of a mucosal adjuvant. Immunology and cell biology 88, 698–706 (2010).

13 Arakawa, T. et al. A plant-based cholera toxin B subunit–insulin fusion protein protects against the development of autoimmune diabetes. Nature biotechnology 16, 934–938 (1998).

14 Tarkowski, A., Sun, J.B., Holmdahl, R., Holmgren, J. & Czerkinsky, C. Treatment of experimental autoimmune arthritis by nasal administration of a type II collagen–cholera toxoid conjugate vaccine. Arthritis & Rheumatism: Official Journal of the American College of Rheumatology 42, 1628–1634 (1999).

15 Rask, C. et al. Prolonged oral treatment with low doses of allergen conjugated to cholera toxin B subunit suppresses immunoglobulin E antibody responses in sensitized mice. Clinical and experimental allergy: journal of the British Society for Allergy and Clinical Immunology 30, 1024–1032 (2000).

16 Phipps, P.A. et al. Prevention of mucosally induced uveitis with a HSP60□derived peptide linked to cholera toxin B subunit. European journal of immunology 33, 224–232 (2003).

17 Ruhlman, T., Ahangari, R., Devine, A., Samsam, M. & Daniell, H. Expression of cholera toxin B–proinsulin fusion protein in lettuce and tobacco chloroplasts–oral administration protects against development of insulitis in non obese diabetic mice. Plant biotechnology journal 5, 495–510 (2007).

18 Sun, J.B., Czerkinsky, C. & Holmgren, J. Mucosally induced immunological tolerance, regulatory T cells and the adjuvant effect by cholera toxin B subunit. Scandinavian journal of immunology 71, 1–11 (2010).

19 Hamorsky, K.T. et al. Rapid and scalable plant-based production of a cholera toxin B subunit variant to aid in mass vaccination against cholera outbreaks. PLoS neglected tropical diseases 7, e2046 (2013).

20 Reeves, M.A. et al. Spray-Dried Formulation of Epicertin, a Recombinant Cholera Toxin B Subunit Variant That Induces Mucosal Healing. Pharmaceutics 13, 576 (2021).

21 Baldauf, K. et al. Oral administration of a recombinant cholera toxin B subunit promotes mucosal healing in the colon. Mucosal immunology 10, 887–900 (2017).

22 Morris, D.A., Reeves, M.A., Royal, J.M., Hamorsky, K.T. & Matoba, N. Isolation and detection of a KDEL-tagged recombinant cholera toxin B subunit from Nicotiana benthamiana. Process Biochem 101, 42–49, doi:10.1016/j.procbio.2020.10.018 (2021).

23 Royal, J.M. et al. A modified cholera toxin B subunit containing an ER retention motif enhances colon epithelial repair via an unfolded protein response. FASEB J 33, 13527–13545, doi:10.1096/fj.201901255R (2019).

24 Royal, J.M., Reeves, M.A. & Matoba, N. Repeated Oral Administration of a KDEL-tagged Recombinant Cholera Toxin B Subunit Effectively Mitigates DSS Colitis Despite a Robust Immunogenic Response. Toxins (Basel) 11, doi:10.3390/toxins11120678 (2019).

25 Tuse, D. et al. Pharmacokinetics and Safety Studies in Rodent Models Support Development of EPICERTIN as a Novel Topical Wound-Healing Biologic for Ulcerative Colitis. J Pharmacol Exp Ther 380, 162–170, doi:10.1124/jpet.121.000904 (2022).

26 Woogen, S.D., Ealding, W. & Elson, C.O. Inhibition of murine lymphocyte proliferation by the B subunit of cholera toxin. The Journal of Immunology 139, 3764–3770 (1987).

27 George-Chandy, A. et al. Cholera toxin B subunit as a carrier molecule promotes antigen presentation and increases CD40 and CD86 expression on antigen-presenting cells. Infection and immunity 69, 5716–5725 (2001).

28 Canziani, G.A., Klakamp, S. & Myszka, D.G. Kinetic screening of antibodies from crude hybridoma samples using Biacore. Anal Biochem 325, 301–307, doi:10.1016/j.ab.2003.11.004 (2004).

29 Matoba, N., Davis, K.R. & Palmer, K.E. Recombinant protein expression in Nicotiana. Methods Mol Biol 701, 199–219, doi:10.1007/9781-61737-957-4_11 (2011).

30 Kohler, G. & Milstein, C. Continuous cultures of fused cells secreting antibody of predefined specificity. Nature 256, 495–497, doi:10.1038/256495a0 (1975).

31 Tabll, A., Abbas, A.T., El-Kafrawy, S. & Wahid, A. Monoclonal antibodies: Principles and applications of immmunodiagnosis and immunotherapy for hepatitis C virus. World J Hepatol 7, 2369–2383, doi:10.4254/wjh.v7.i22.2369 (2015).

32 Yu, R.K., Usuki, S., Itokazu, Y. & Wu, H.C. Novel GM1 ganglioside-like peptide mimics prevent the association of cholera toxin to human intestinal epithelial cells in vitro. Glycobiology 26, 63–73, doi:10.1093/glycob/cwv080 (2016).

33 Royal, J.M. et al. A modified cholera toxin B subunit containing an ER retention motif enhances colon epithelial repair via an unfolded protein response. The FASEB Journal 33, 13527–13545 (2019).

34 Holmner, A., Askarieh, G., Okvist, M. & Krengel, U. Blood group antigen recognition by Escherichia coli heat-labile enterotoxin. J Mol Biol 371, 754–764, doi:10.1016/j.jmb.2007.05.064 (2007).

35 Lebens, M. et al. Synthesis of hybrid molecules between heat-labile enterotoxin and cholera toxin B subunits: potential for use in a broad-spectrum vaccine. Infect Immun 64, 2144–2150, doi:10.1128/iai.64.6.2144-2150.1996 (1996).

36 Jobling, M.G. & Holmes, R.K. Mutational analysis of ganglioside GM1-binding ability, pentamer formation, and epitopes of cholera toxin B (CTB) subunits and CTB/heat-labile enterotoxin B subunit chimeras. Infection and immunity 70, 1260–1271 (2002).

37 Holmes, R.K. & Twiddy, E.M. Characterization of monoclonal antibodies that react with unique and cross-reacting determinants of cholera enterotoxin and its subunits. Infection and immunity 42, 914–923 (1983).

38 Robb, M., Nichols, J.C., Whoriskey, S.K. & Murphy, J.R. Isolation of hybridoma cell lines and characterization of monoclonal antibodies against cholera enterotoxin and its subunits. Infect Immun 38, 267–272, doi:10.1128/iai.38.1.267-272.1982 (1982).

39 Chou, S.F. Production and purification of monoclonal and polyclonal antibodies against cholera toxin. Hybrid Hybridomics 23, 258–261, doi:10.1089/1536859041651376 (2004).

40 Kenimer, J.G., Probst, P.G., Karpas, A.B., Burns, D.L. & Kaslow, H.R. Monoclonal antibodies against the enzymatic subunit of both pertussis and cholera toxins. Dev Biol Stand 73, 133–141 (1991).

41 Remmers, E.F., Colwell, R.R. & Goldsby, R.A. Production and characterization of monoclonal antibodies to cholera toxin. Infect Immun 37, 70–76, doi:10.1128/iai.37.1.70-76.1982 (1982).

42 Ludwig, D.S., Holmes, R.K. & Schoolnik, G.K. Chemical and immunochemical studies on the receptor binding domain of cholera toxin B subunit. J Biol Chem 260, 12528–12534 (1985).

43 Otte, L., Knaute, T., Schneider-Mergener, J. & Kramer, A. Molecular basis for the binding polyspecificity of an anti-cholera toxin peptide 3 monoclonal antibody. J Mol Recognit 19, 49–59, doi:10.1002/jmr.757 (2006).

44 Anglister, J. & Zilber, B. Antibodies against a peptide of cholera toxin differing in cross-reactivity with the toxin differ in their specific interactions with the peptide as observed by 1H NMR spectroscopy. Biochemistry 29, 921–928, doi:10.1021/bi00456a011 (1990).

45 Tinker, J.K., Erbe, J.L., Hol, W.G. & Holmes, R.K. Cholera holotoxin assembly requires a hydrophobic domain at the A-B5 interface: mutational analysis and development of an in vitro assembly system. Infect Immun 71, 4093–4101, doi:10.1128/IAI.71.7.4093-4101.2003 (2003).

46 Reichow, S.L., Korotkov, K.V., Hol, W.G. & Gonen, T. Structure of the cholera toxin secretion channel in its closed state. Nat Struct Mol Biol 17, 1226–1232, doi:10.1038/nsmb.1910 (2010).

47 Torgersen, M.L., Skretting, G., van Deurs, B. & Sandvig, K. Internalization of cholera toxin by different endocytic mechanisms. J Cell Sci 114, 3737–3747 (2001).

48 Belisle, B.W., Twiddy, E.M. & Holmes, R.K. Monoclonal antibodies with an expanded repertoire of specificities and potent neutralizing activity for Escherichia coli heat-labile enterotoxin. Infect Immun 46, 759–764, doi:10.1128/iai.46.3.759-764.1984 (1984).

49 Sethi, A. et al. Cell type and receptor identity regulate cholera toxin subunit B (CTB) internalization. Interface Focus 9, 20180076, doi:10.1098/rsfs.2018.0076 (2019).

50 Wands, A.M. et al. Fucosylation and protein glycosylation create functional receptors for cholera toxin. Elife 4, e09545, doi:10.7554/eLife.09545 (2015).

51 Cervin, J. et al. GM1 ganglioside-independent intoxication by Cholera toxin. PLoS Pathog 14, e1006862, doi:10.1371/journal.ppat.1006862 (2018).

52 Jia, J. et al. KDEL receptor is a cell surface receptor that cycles between the plasma membrane and the Golgi via clathrin-mediated transport carriers. Cell Mol Life Sci 78, 1085–1100, doi:10.1007/s00018-020-03570-3 (2021).

53 Kozireski-Chuback, D., Wu, G. & Ledeen, R.W. Developmental appearance of nuclear GM1 in neurons of the central and peripheral nervous systems. Developmental brain research 115, 201–208 (1999).

54 Okada, E., Maeda, T. & Watanabe, T. Immunocytochemical study on cholera toxin binding sites by monoclonal anti-cholera toxin antibody in neuronal tissue culture. Brain research 242, 233–241 (1982).

55 Cervin, J. et al. GM1 ganglioside-independent intoxication by Cholera toxin. PLoS pathogens 14, e1006862 (2018).

56 Moreno Altamirano, M.M.B., Aguilar Carmona, I. & Sánchez□Garcia, F.J. Expression of GM1, a marker of lipid rafts, defines two subsets of human monocytes with differential endocytic capacity and lipopolysaccharide responsiveness. Immunology 120, 536–543 (2007).

57 Francis, M. et al. Cyclic AMP-independent effects of cholera toxin on B cell activation. II. Binding of ganglioside GM1 induces B cell activation. The Journal of Immunology 148, 1999–2005 (1992).

58 Smits, H. et al. Cholera toxin B suppresses allergic inflammation through induction of secretory IgA. Mucosal immunology 2, 331–339 (2009).

59 Holmgren, J., Lönnroth, I. & Svennerholm, L. Tissue receptor for cholera exotoxin: postulated structure from studies with GM1 ganglioside and related glycolipids. Infection and immunity 8, 208–214 (1973).

60 Nystrom, E.E.L. et al. An intercrypt subpopulation of goblet cells is essential for colonic mucus barrier function. Science 372, doi:10.1126/science.abb1590 (2021).

